# Novel markers for haemogenic endothelium and haematopoietic progenitors in the mouse yolk sac

**DOI:** 10.1101/2025.11.21.689693

**Authors:** Guillermo Diez-Pinel, Alessandro Muratore, Christiana Ruhrberg, Giovanni Canu

## Abstract

Erythro-myeloid progenitors (EMPs) originate from the haemogenic endothelium in the yolk sac via an endothelial-to-haematopoietic transition (EHT) to generate blood and immune cells that support embryo development. Yet, the transitory nature of EHT and the limited availability of molecular markers has constrained our understanding of the origin, identity and differentiation dynamics of EMPs. Here, we have refined the annotation of yolk sac haemato-vascular populations in publicly available single-cell RNA sequencing (scRNAseq) datasets from mouse embryos to identify novel molecular markers of haemogenic endothelium and EMPs. By sub-clustering key cell populations followed by pseudotime analysis, we refined cluster annotations and then reconstructed differentiation trajectories. Subsequent differential gene expression analysis between clusters identified novel cell surface markers for haemogenic endothelial cells (*Fxyd5* and *Scarf1*) and EMPs (*Fcer1g*, *Tyrobp* and *Mctp1*). Further, we have identified candidate signalling and metabolic pathways that may regulate yolk sac haematopoietic emergence and differentiation. The specificity of FXYD5, SCARF1 and FCER1G for haemogenic endothelium and EMPs was validated by immunostaining of mouse yolk sac. These insights into the transcriptional dynamics in the yolk sac should support future investigation of EHT and haematopoietic differentiation during early mammalian development.

## INTRODUCTION

Haematopoiesis in the developing embryo proceeds in three temporally and spatially overlapping waves in close proximity to vascular endothelial cells (Ivanovs *et al*., 2017; Canu and Ruhrberg, 2021). The first wave, known as primitive haematopoiesis, arises in the yolk sac, when extra-embryonic mesoderm differentiates into vascular and haematopoietic progenitors that progressively organise into blood islands, and produces nucleated erythrocytes, megakaryocytes and macrophages (Ferkowicz and Yoder, 2005). This process occurs in the mouse from embryonic day (E) 7.0 onwards, equivalent to 2-3 post-conception weeks in human.

The second haematopoietic wave, known as pro-definitive or transient definitive haematopoiesis, also emerges in the yolk sac, when a change in cellular identity causes haematopoietic progenitors to bud off the endothelium as clusters of round cells, a process called endothelial-to-haematopoietic transition (EHT) (Frame *et al*., 2016). In the mouse yolk sac, EHT generates mainly erythro-myeloid progenitors (EMPs) from E8.5, alongside lympho-myeloid progenitors (LMPs) from E9.5 (Böiers *et al*., 2013; Frame, McGrath and Palis, 2013). Once the blood circulation is established by E9.5, these progenitors are transported out of the yolk sac and seed the foetal liver, from where they contribute blood and immune cells from mid-gestation until birth (Hoeffel *et al*., 2015). Additionally, we recently showed that haematopoietic progenitors akin to EMPs can also arise from a *Pax3*-expressing lineage in the mouse and transiently seed the foetal liver by E12.5 to contribute erythroid and myeloid cells to foetal haematopoiesis (Canu *et al*., 2024). Based on their origin from a *Pax3*-expressing progenitor, these haematopoietic cells may arise via EHT from a paraxial mesoderm-derived haemogenic endothelium, although further work is needed to address this possibility. Among the early haematopoietic populations, yolk sac-derived EMPs generate blood and immune cells important for embryogenesis (Frame, McGrath and Palis, 2013; Hoeffel *et al*., 2015) and self-renewing tissue-resident macrophages that persist into adulthood (Moskalik, Niderla-Bielińska and Ratajska, 2021; Jiang *et al*., 2024; Liu *et al*., 2024).

The third haematopoietic wave, known as definitive haematopoiesis, originates intra-embryonically in the dorsal aorta, where EHT produces haematopoietic stem cells (HSCs) that enter the circulation to first colonise the foetal liver by mid-gestation and later seed the bone marrow to establish long-term, self-renewing adult haematopoiesis (Tavian *et al*., 1996; Oberlin *et al*., 2010; Ivanovs *et al*., 2011; Canu and Ruhrberg, 2021; Sánchez-Lanzas, Jiménez-Pompa and Ganuza, 2024; Sugimura *et al*., 2025). HSCs are first produced at E10.5 in the mouse, and 4-5 post-conception weeks in human.

Despite their key roles in embryonic haematopoiesis and the generation of tissue macrophages with a role in adult homeostasis, the transitory nature of EMPs and their overlapping markers with HSC lineages have made the study of EMP-derived cell lineages challenging. For example, the Lin^-^ KIT^+^ CD41^low^ CD16/32^+^ cell surface profile allows EMP isolation by flow cytometry from the early embryo, but is insufficient to unambiguously distinguish EMPs from HSC-derived myeloid progenitors at mid-gestation (McGrath *et al*., 2015; Van Deren *et al*., 2022). Temporally-controlled lineage tracing via Cre-mediated reporter gene activation has therefore been used as a complementary approach to distinguish EMPs from HSC-derived progenitors based on the earlier production of EMPs (Gomez Perdiguero *et al*., 2014; Hoeffel *et al*., 2015; McGrath, Frame and Palis, 2016). However, this approach relies on transgenic mice expressing suitable Cre drivers and expression reporters. Therefore, specific molecular markers are needed to allow accurate EMP isolation and to study their formation and behaviour in the mouse embryo. Moreover, such markers might facilitate the identification of analogous markers suitable to study foetal haematopoiesis in human embryos, in which genetic lineage tracing is not an option.

Here, we have leveraged publicly available mouse embryonic single-cell RNA sequencing (scRNAseq) data to investigate cell populations in the developing mouse yolk sac to identify novel candidate markers for haemogenic endothelium and EMPs.

## METHODS

### Integration of publicly available E8.5 scRNAseq mouse embryo data

The file ‘embryo_sce.rds’ containing a processed SingleCellExperiment object including scRNAseq data from the ArrayExpress dataset <u>E-MTAB-6967</u> (Pijuan-Sala *et al*., 2019) and ArrayExpress dataset <u>E-MTAB-11763</u> (Goh *et al*., 2023) was downloaded from https://marionilab.github.io/ExtendedMouseAtlas/. The cells annotated as embryonic day E8.5 were subset for analysis in R v4.5.1, with Rstudio v2025.05.1 using the Seurat v4.5.1 R package (Satija *et al*., 2015). The function ‘as.Seurat’ was used to generate a Seurat object from the SingleCellExperiment object. For sample integration, the E-MTAB-11763 cells were subset and split into layers containing individual samples, counts were log-normalised with the ‘NormalizeData’ function, the top-2000 variable features were identified with the ‘FindVariableFeatures’ and scaled and centred with the ‘ScaleData’ function. Then, PCA dimensionality reduction was performed with the ‘RunPCA’ function and Harmony Integration (Korsunsky *et al*., 2019) was performed using the ‘IntegrateLayers’ function. A Uniform Manifold Approximation and Projection (UMAP) reduction was obtained with the ‘RunUMAP’ function, for which the number of principal components was selected by examination of an elbow plot. Lastly, the layers were joined with the ‘JoinLayers’ function.

This same integration pipeline was used for all other data integration in the study.

### Re-annotation of yolk sac cell types in E8.5 scRNAseq data

Cells with anatomy ‘Yolk sac’ (YS) from haemato-vascular cell types were subset and integrated by sample. For clustering, a k-nearest neighbour (KNN) graph was calculated (k = 20), and the Louvain algorithm was used for graph-clustering. Then, the expression of known cell type markers in the yolk sac clusters was used alongside the anatomical origin (yolk sac or embryo) to annotate these haemato-vascular yolk sac cells as well as the embryo haemato-vascular cells more accurately. Afterwards, both yolk sac and embryo cells were merged and integrated by sample. These data were then used as the reference dataset for reference-based annotation of the E-MTAB-6967 haemato-vascular cells using the SingleR v2.10.0 R package (Aran *et al*., 2019), and then merged and integrated by sample with the E-MTAB-11763 data.

### Analysis of gene expression along the EHT differentiation trajectory

To identify haemogenic endothelium cells, we sub-clustered all cells annotated as endothelial cells and EMPs in the combined dataset, including Allantois endothelium, Venous endothelium, Embryo proper endothelium, Endocardium, YS endothelium, Vitellin vein endothelium, EMPs and EMP-like embryo, and then examined the expression of haemato-vascular markers.To analyse the EHT differentiation trajectory, we sub-clustered yolk sac haemato-vascular cells, including YS endothelium, haemogenic endothelium, EMPs, BP1, BP2 and Erythroid, and the resulting UMAP reduction was used to carry out a pseudotime analysis to predict differentiation trajectories with the Monocle3 v1.4.26 R package (Cao *et al*., 2019). Genes that changed as a function of pseudotime were identified with the ‘graph_test’ function and grouped according to their expression patterns into modules with the ‘find_gene_modules’, which were then grouped manually into four supermodules (downwards trend, upwards trend, peaking at EMPs and peaking at BP1).

### Gene ontology and differential gene expression of haemato-vascular clusters

Enrichment analyses against the GeneOntology: Biological Pathway and WikiPathways databases were carried out using the ‘enrichGO’ and ‘enrichWP’ functions from the clusterProfiler v4.16.0 R package (Wu *et al*., 2021), respectively. All genes with at least 1 count were used as the background universe. Cytoscape 3.10.3 (Shannon *et al*., 2003) and the RCy3 v2.28.1 R package (Gustavsen *et al*., 2019) were used to plot the TYROBP WikiPathway pathway. For surface marker identification, differential expression analyses were carried out with the ‘FindMarkers’ function, with an adjusted P-value threshold of 0.05. Then, Uniprot accession IDs were retrieved from the org.Mm.eg.db v3.21.0 R package (Carlson, 2025) using the ‘select’ function from the AnnotationDbi v1.70.0 R package (Pagès *et al*., 2025). A list of curated transmembrane proteins was obtained from Uniprot (https://www.uniprot.org/) on the 30th of July 2025 using the query ‘(keyword:KW-0812) AND (reviewed:true)’, and filtering for ‘Mouse’ as the relevant organism. This list was then used to filter the differential expression results for transmembrane proteins. Plotting of UMAP reductions was carried out with Seurat’s ‘DimPlot’ and the function ‘DimPlot_scCustom’ from the scCustomize v3.2.0 package (Marsh *et al*., 2025). Plotting of normalised gene expression onto UMAP reductions was carried out with Seurat’s ‘FeaturePlot’ and scCustomize’s ‘FeaturePlot_scCustom’ functions. Other plots were generated using the ggplot2 v4.0.0 R package (Wickham *et al*., 2025).

### Animal procedures and tissue staining

Animal procedures were performed with Animal Welfare Ethical Review Body (AWERB) and UK Home Office approval. C57BL/6J mice were timed-mated for embryo collection at E8.5 after cervical dislocation culling of the pregnant dam. Yolk sacs were fixed in 4% formaldehyde, washed in phosphate-buffered saline (PBS) and then blocked in PBS containing 2% serum-free protein block (DAKO), 2% bovine serum albumin and 0.4% Triton X-100 before staining with the following antibodies: goat anti-mouse KIT (R&D Systems #AF1356-SP, 1:50), rat anti-mouse PECAM1 (BD Pharmingen #553370, 1:50), rabbit anti-mouse FXYD5/Dysadherin (Proteintech #12166-1-AP, 1:50), rabbit anti-mouse SCARF1 (Proteintech #13702-1-AP, 1:100) or rabbit anti-mouse FCER1G (GeneTex #GTX108487s, 1:50), followed by the appropriate secondary antibodies, which were Alexa Fluor 488-conjugated donkey anti-goat Fab fragments (Stratech #705-547-003, 1:200), Cy3-conjugated donkey anti-rat Fab fragments (Stratech #712-166-150, 1:200), and Alexa Fluor 647-conjugated donkey anti-rabbit Fab fragments (Stratech #711-607-003, 1:200). Secondary-only controls were included but are not shown. DAPI-counterstained sections were imaged on a LSM710 confocal microscope (Zeiss).

## RESULTS

### Refined annotation of E8.5 yolk sac and embryo cell clusters

To identify the transcriptomic signature of yolk sac haemogenic endothelium and that of its haemopoietic progeny, we re-analysed published scRNAseq data from E8.5 mouse embryos (**Fig. 1A**), which were extracted from the ArrayExpress datasets E-MTAB-6967 (Pijuan-Sala *et al*., 2019) and E-MTAB-11763 (Goh *et al*., 2023). E-MTAB-6967 contains data from E6.5 to E8.5 mouse embryos, including yolk sac and embryo proper, which were sequenced and analysed together to construct a molecular atlas describing early mouse gastrulation (Pijuan-Sala *et al*., 2019). E-MTAB-11763 instead contains data that were generated by collecting yolk sac and embryo proper from E8.5 to E9.5 mouse embryos and sequencing them separately. These two published datasets overall comprise 400,000 cells from E6.5 to E9.5. A subsequent *in silico* merge of E-MTAB-11763 data and E-MTAB-6967 embryo and yolk sac data was previously analysed to compare the mouse versus human yolk sac contribution to metabolic and nutritional support as well as to early erythroid and myeloid lineages (Goh *et al*., 2023). As the goal of our analysis was to define the transcriptome of yolk sac endothelium and its haematopoietic progeny at the time when EMPs emerge from yolk sac haemogenic endothelium, we subset only the E8.5 data from E-MTAB-6967 and E-MTAB-11763 (**Fig. 1A**).

**Fig 1.**
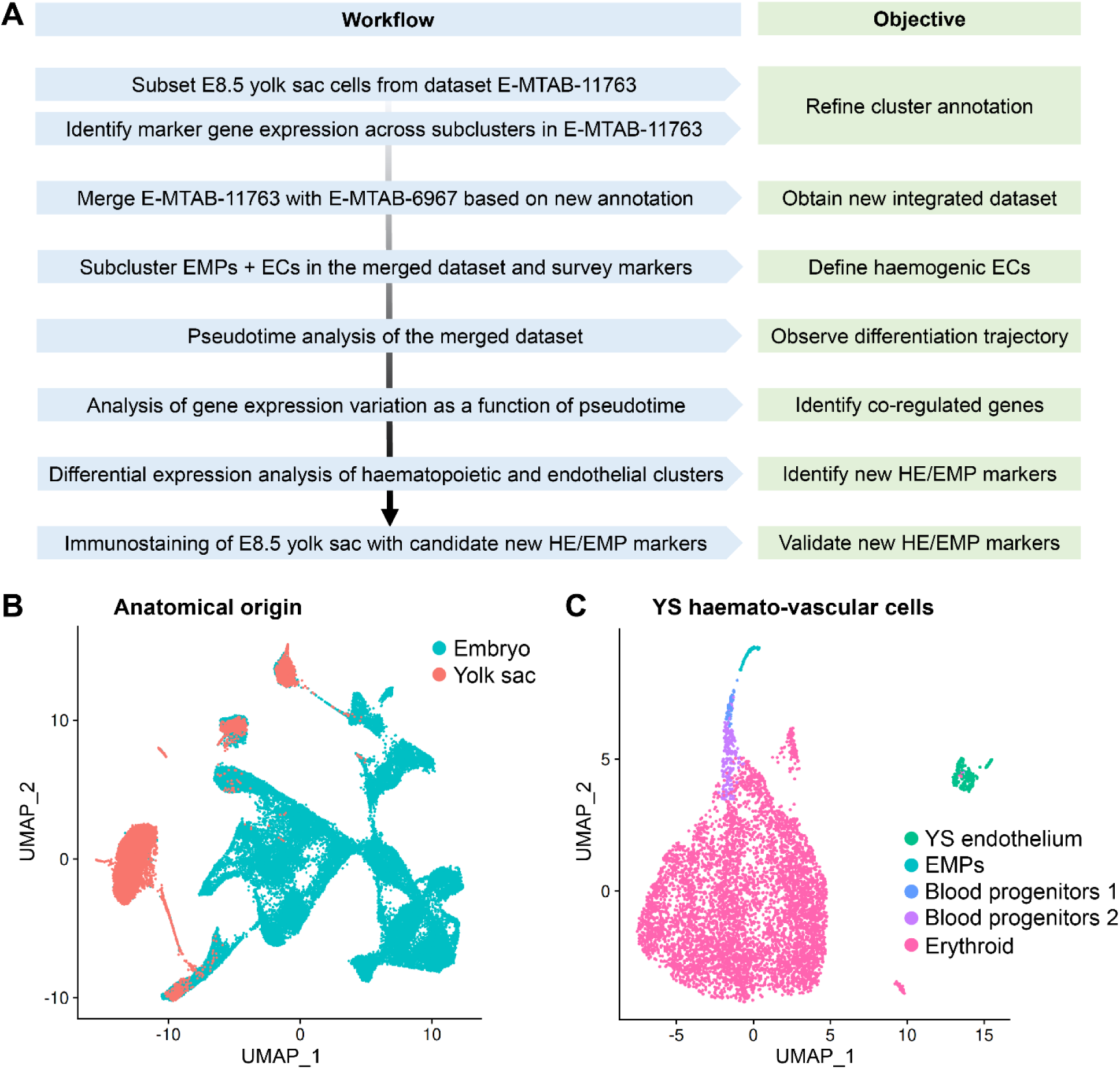
Re-analysis of E8.5 mouse embryonic scRNAseq data to study yolk sac EHT. **A** Workflow for the integrated re-analysis of mouse embryonic scRNAseq datasets E-MTAB-11763 and E-MTAB-6967. **B** UMAP of dataset E-MTAB-11763 showing cells sequenced from the E8.5 mouse embryo and labelled according to their anatomical origin as embryo and yolk sac. **C** UMAP obtained by subsetting haemato-vascular cells from E8.5 yolk sac from E-MTAB-11763 and labelled based on our refined cluster annotation. EMP: erythro-myeloid progenitor; EC: endothelial cell; HE: haemogenic endothelium; YS: yolk sac.

First, we analysed E8.5 data from E-MTAB-11763, which contained yolk sac and embryo proper data annotated separately, and generated a UMAP including 75 clusters, using the cluster annotations provided by the authors of the original publication (**Fig. S1**). Next, we selectively sub-clustered the 8,453 haemato-vascular cells that had been sequenced from the yolk sac (**Fig. 1B,C**). To confirm the identity of the resulting clusters, we analysed the expression of known markers of endothelial and haematopoietic progenitor populations (**Fig. 2A-D**). The yolk sac endothelial cell (EC) cluster was identified via expression of the pan-EC markers *Kdr* and *Cdh5* together with *Lyve1* and *Stab2* (**Fig. 2A**) as two markers typical for yolk sac ECs (Gordon, Gale and Harvey, 2008; Shvartsman *et al*., 2019). The EMP cluster was identified via detection of *Csf1r* and *Ptprc* as well as the myeloid markers *Spi1 and Mpo* (**Fig. 2B**). Notably, *Kdr*, *Cdh5* and *Lyve1* were also detected in the EMP cluster (**Fig. 2A**), agreeing with their known endothelial origin (Gomez Perdiguero *et al*., 2014). Clusters expressing haematopoietic progenitor markers (*Vav1*, *Myb*, *Gata2*) (**Fig. 2C**) together with erythroid (*Gata1*, *Klf1*) or myeloid (*Spi1, Mpo)* differentiation markers (**Fig. 2D**) represented intermediate progenitors at different states of commitment and were annotated as blood progenitors (BP) 1 and BP2 (**Fig. 1C**). Finally, high expression levels of differentiation markers (*Alas2*, *Gypa*) (**Fig. 2D**), in the absence of endothelial (*Cdh5*, *Lyve1*), EMP (*Csf1r*, *Ptprc*) or progenitor markers (*Spi1*, *Gata2*), identified a cluster of differentiated erythroid cells.

**Fig 2.**
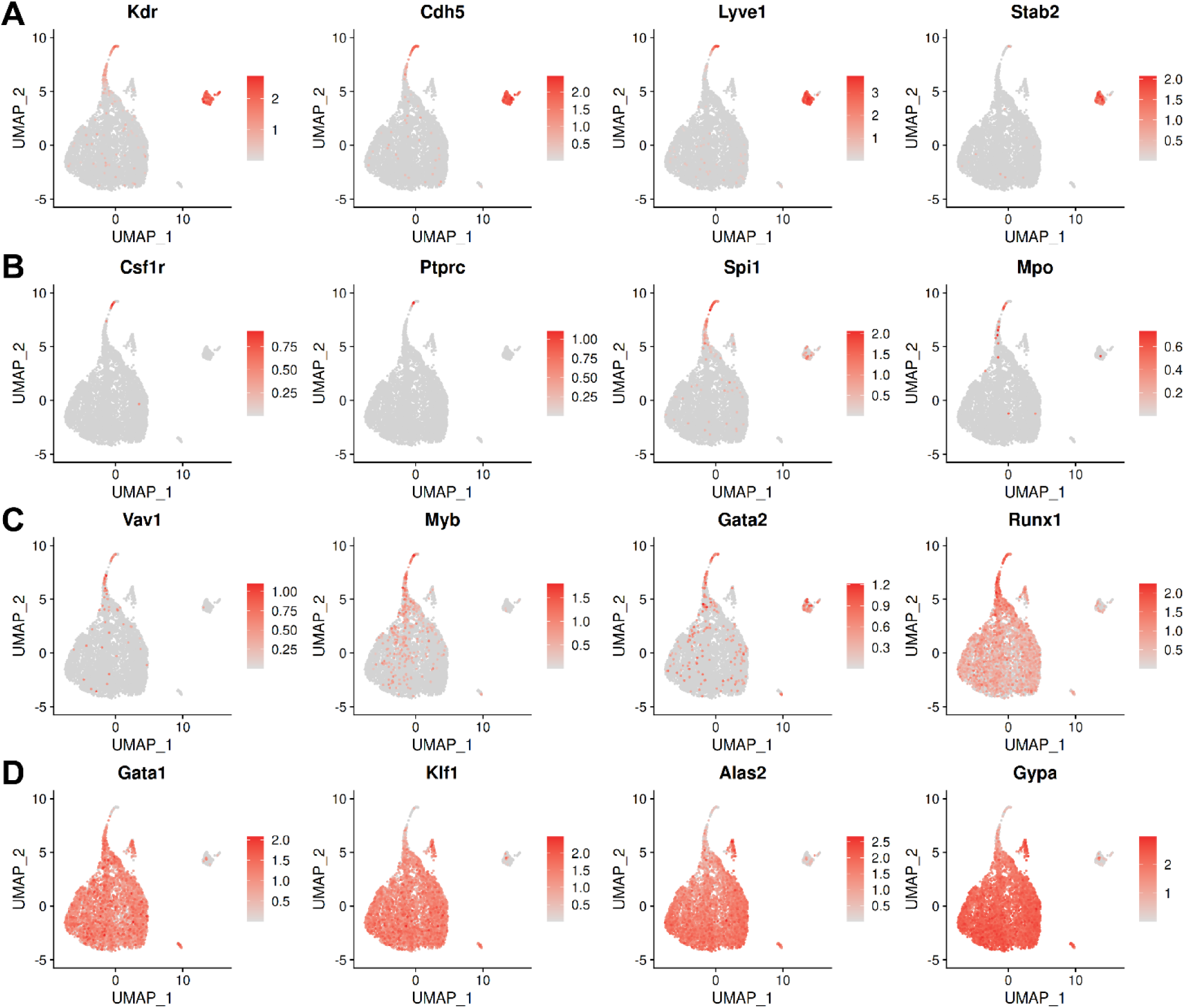
Marker analysis identifies haemato-vascular cell populations in E8.5 yolk sac data in E-MTAB-11763. UMAP representation of transcript levels for the indicated marker genes for (**A**) endothelial cells, (**B**) EMPs, (**C**) haematopoietic progenitors and (**D**) erythroid cells. The scale bar represents log-normalised transcript levels.

As expected, the detection levels of endothelial and haematopoietic genes displayed opposite trends between the endothelial and haematopoietic clusters. Thus, *Kdr* and *Cdh5* appeared highly expressed in ECs and EMPs but their detection declined in BP1, whilst they were barely detectable in BP2 and absent from erythroid cells (**Fig. 2A**). Conversely, erythroid genes such as *Klf1* and *Gypa* were not detected in ECs and EMPs, detected at low levels in BP1, at higher levels in BP2 and at very high levels in erythroid cells (**Fig. 2D**). Known haematopoietic progenitor markers (*e.g.*, *Myb*, *Gata2, Runx1*) displayed a gradual decrease from EMPs to BP1 and BP2, supporting the notion that the three clusters represented successive maturation stages (**Fig. 2C**). Myeloid markers (*e.g.*, *Csf1r, Spi1, Mpo*) did not identify a separate cluster but were present in the EMP cluster and, at lower levels, also in BP1 (**Fig. 2B**), agreeing with prior studies suggesting that macrophages are only present in the yolk sac from E9.0 onwards (Hoeffel and Ginhoux, 2018).

Unexpectedly, we observed that E-MTAB-11763 contained cells that had been isolated and sequenced from the embryo proper but nevertheless clustered with yolk sac EMPs and BPs; accordingly, we annotated these cells as EMP-like embryo and blood progenitors embryo, respectively (**Fig. 3A,B**). Also unexpectedly, some cells sequenced from the embryo proper had been annotated in the original publication describing E-MTAB-11763 as representing yolk sac-derived ECs. As the endothelium in the yolk sac vasculature forms a continuum with the endothelium of the vitelline vein, cells within the embryo proper that resemble yolk sac ECs likely represent vitelline vein ECs, and we re-annotated them accordingly (**Fig. 3A,B**).

**Fig 3.**
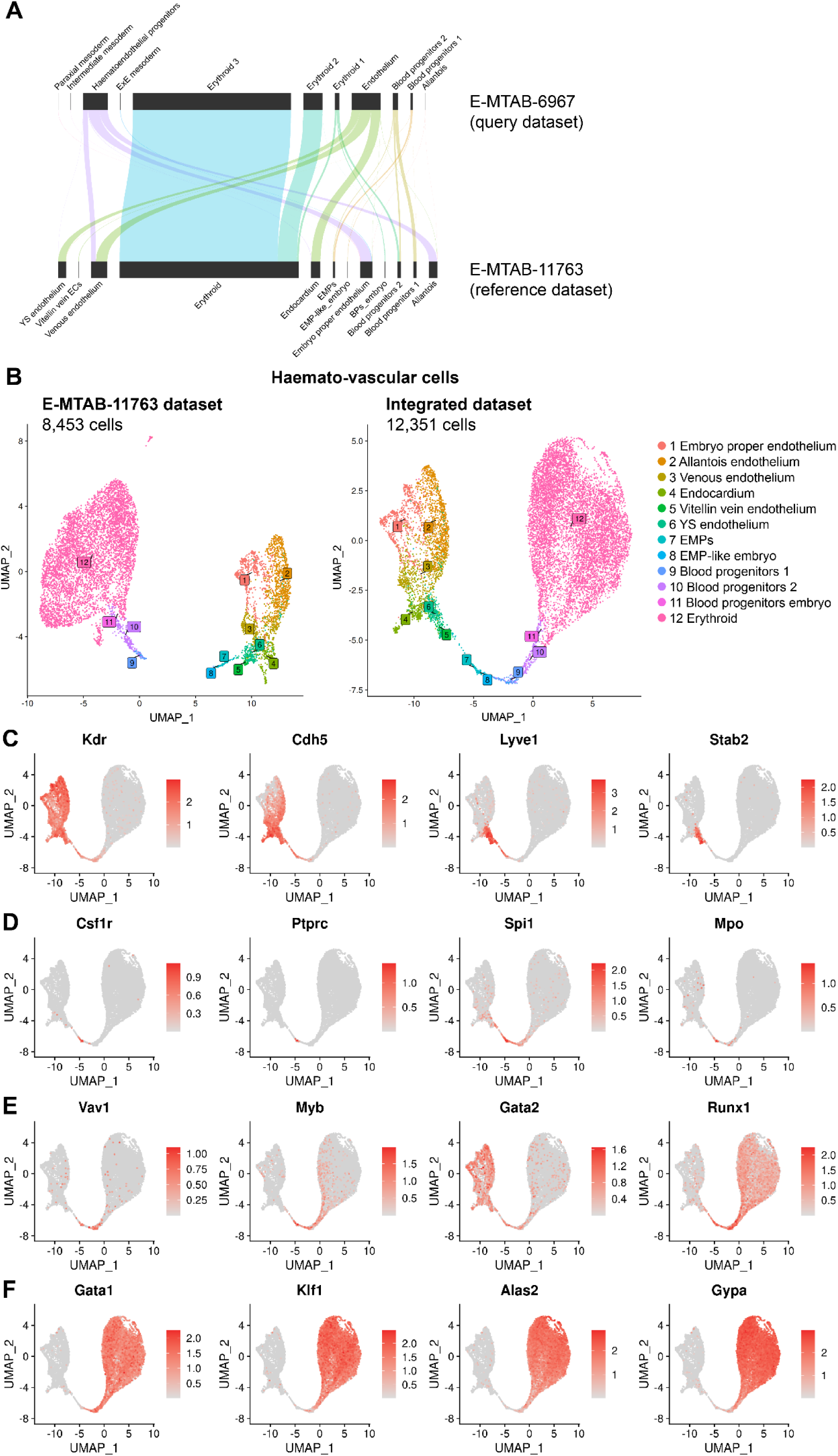
Refined cluster annotation of E8.5 embryo and yolk sac haemato-vascular cell clusters after integration of E-MTAB-11763 and E-MTAB-6967. **A** Reference-based annotation based on gene expression was used to apply our revised annotation for E8.5 data in E-MTAB-11763 to the E8.5 data in E-MTAB-6967 before integrating the two datasets. **B** UMAPs showing the E8.5 haemato-vascular clusters from both embryo and yolk sac for dataset E-MTAB-11763 before integration to dataset E-MTAB-6967 and after reference-mapping and integration of the two datasets. **C-F** UMAP representation of transcript levels for the indicated marker genes for (**C**) endothelial cells, (**D**) EMPs, (**E**) haematopoietic progenitors and (**F**) erythroid cells. The scale bar represents log-normalised transcript levels.

Next, we integrated E-MTAB-11763 with E-MTAB-6967 to increase the number of cells available for downstream analysis in the haemato-vascular clusters to identify yolk sac haemogenic endothelium. As cells derived from the yolk sac and embryo proper had not been separately sequenced to generate E-MTAB-6967, we used a validated, reference-based annotation method (Aran *et al*., 2019) to harmonise cell type annotations and distinguish cells from these sources before integrating the two datasets (**Fig. 3A**). Specifically, this method uses gene expression information to assign the most likely cluster from a reference dataset to a query dataset, which in our case involved applying our revised annotation for E8.5 data in E-MTAB-11763 to the E8.5 data in E-MTAB-6967. This approach increased the overall number of haemato-vascular cells from 8,453 to 12,351 for further analysis (**Fig. 3B**). We also surveyed the expression of key known marker genes of the cell populations of interest to confirm the appropriate annotation of the integrated dataset (**Fig. 3C-F**).

### Identifying yolk sac haemogenic ECs via their transcriptomic signature

The haemogenic endothelium is a rare and transient population, and thus difficult to study (Gritz and Hirschi, 2016; Scarfò *et al*., 2024). To identify haemogenic ECs within the yolk sac endothelial cluster, we subset EMPs and ECs from both yolk sac and embryo proper and re-clustered them to generate a new UMAP including only these cells, which was annotated according to our refined nomenclature. Using this approach, yolk sac ECs clustered in an intermediate position between embryo-derived ECs and EMPs and also split into two sub-clusters, one positioned closer to EMPs and the other closer to endocardial ECs (**Fig. 4A**). The analysis of known marker genes showed that both sub-clusters expressed markers of yolk sac endothelium at similar levels (*Lyve1*, *Stab2*), whereas only the EMP-proximal sub-cluster co-expressed haematopoietic progenitor genes (*Runx1*, *Myb*, *Spi1*) (**Fig. 4B**). Together, these findings indicate that a subset of yolk sac ECs have a dual endothelial and haematopoietic cell identity, consistent with a presumed identify of haemogenic ECs that give rise to EMPs.

**Fig 4.**
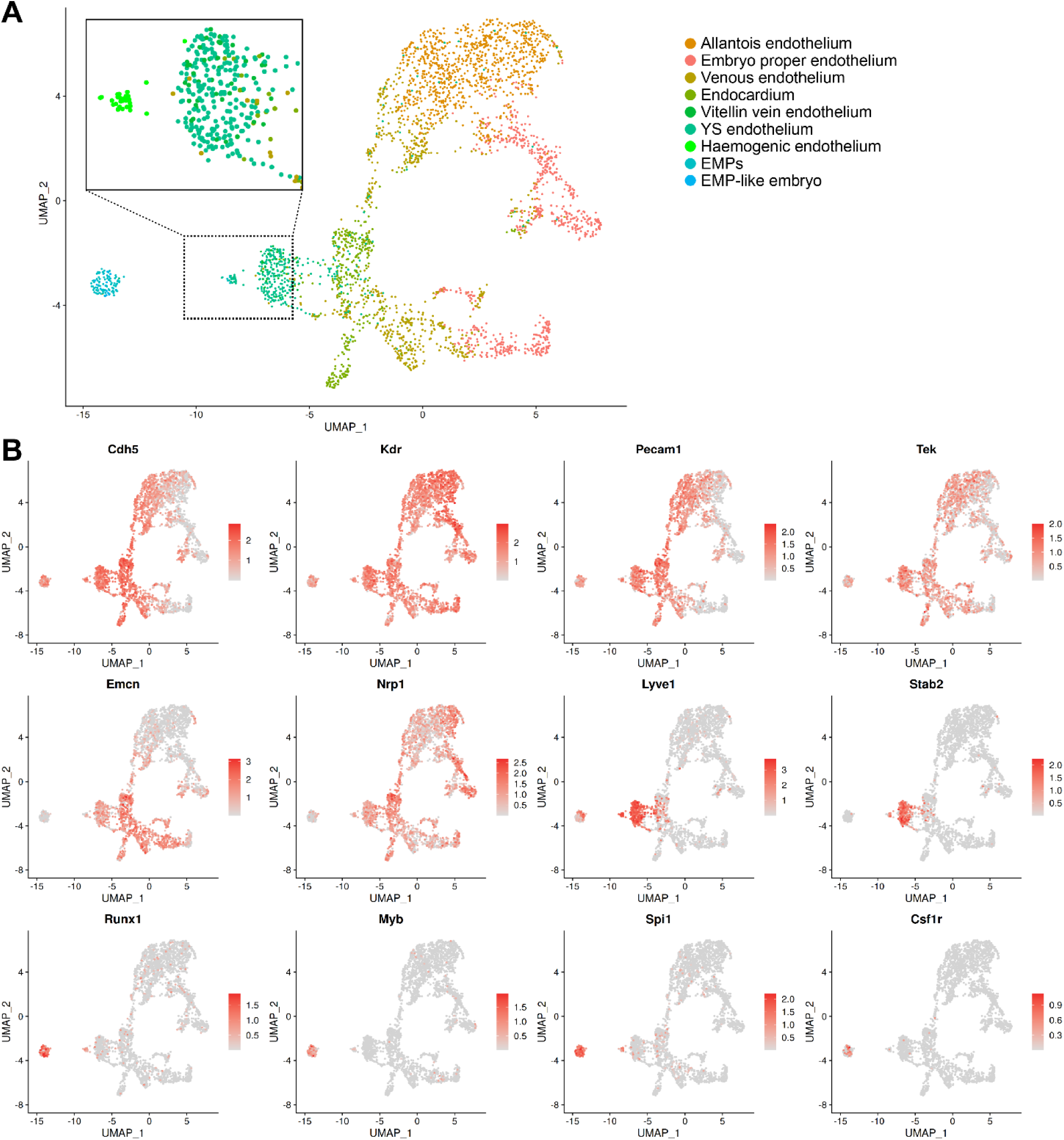
Endothelial cell and EMP sub-clustering identifies haemogenic endothelium. **A** UMAP obtained after subsetting and re-clustering all cells annotated as endothelial cells and EMPs after integration of E8.5 data in E-MTAB-11763 and E-MTAB-6967. The dotted box indicates an area that is zoomed on the cells previously annotated as yolk sac endothelial cells, vitellin vein endothelial cells and EMPs; note the formation of a separate cluster positioned between EMPs and the main yolk sac endothelial cluster consistent with a haemogenic endothelial identify. **B** UMAP representation of key endothelial and haematopoietic markers corroborates the identification of the haemogenic endothelial cluster. The scale bars represent log-normalised transcript levels. YS: yolk sac; EMPs: erythro-myeloid progenitors.

### Pseudotime analysis identifies transcriptomic changes accompanying EHT

Having identified a cell cluster with haemogenic endothelial identity, we next used our refined annotation of the yolk sac populations to study signalling pathways and markers defining the transition from haemogenic ECs into EMPs and BPs. When analysing samples that contain cells along a continuous differentiation process, single-cell transcriptomic data can be used to arrange cells along a pseudotime trajectory (Cannoodt, Saelens and Saeys, 2016). In essence, pseudotime analysis uses gene expression patterns to reorder cells and arrange them based on their presumed progressive gene expression differences, thus reconstructing trajectories along which cells may be differentiating. Thus, we used pseudotime analysis of yolk sac cell populations to study the transition of ECs into haematopoietic populations. In agreement with prior knowledge of EHT, pseudotime analysis of our integrated E8.5 dataset suggested a continuum of cell differentiation from endothelial to haemogenic endothelium and then to haematopoietic cells (**Fig. 5A**), and also highlighted progressive erythroid maturation (**Fig. 5B**). A refined pseudotime analysis after exclusion of erythroid cells as the predominant cell type corroborated the expected progressive transition of yolk sac ECs to haemogenic ECs, EMPs, BP1 and BP2 (**Fig. 5C**).

**Fig 5.**
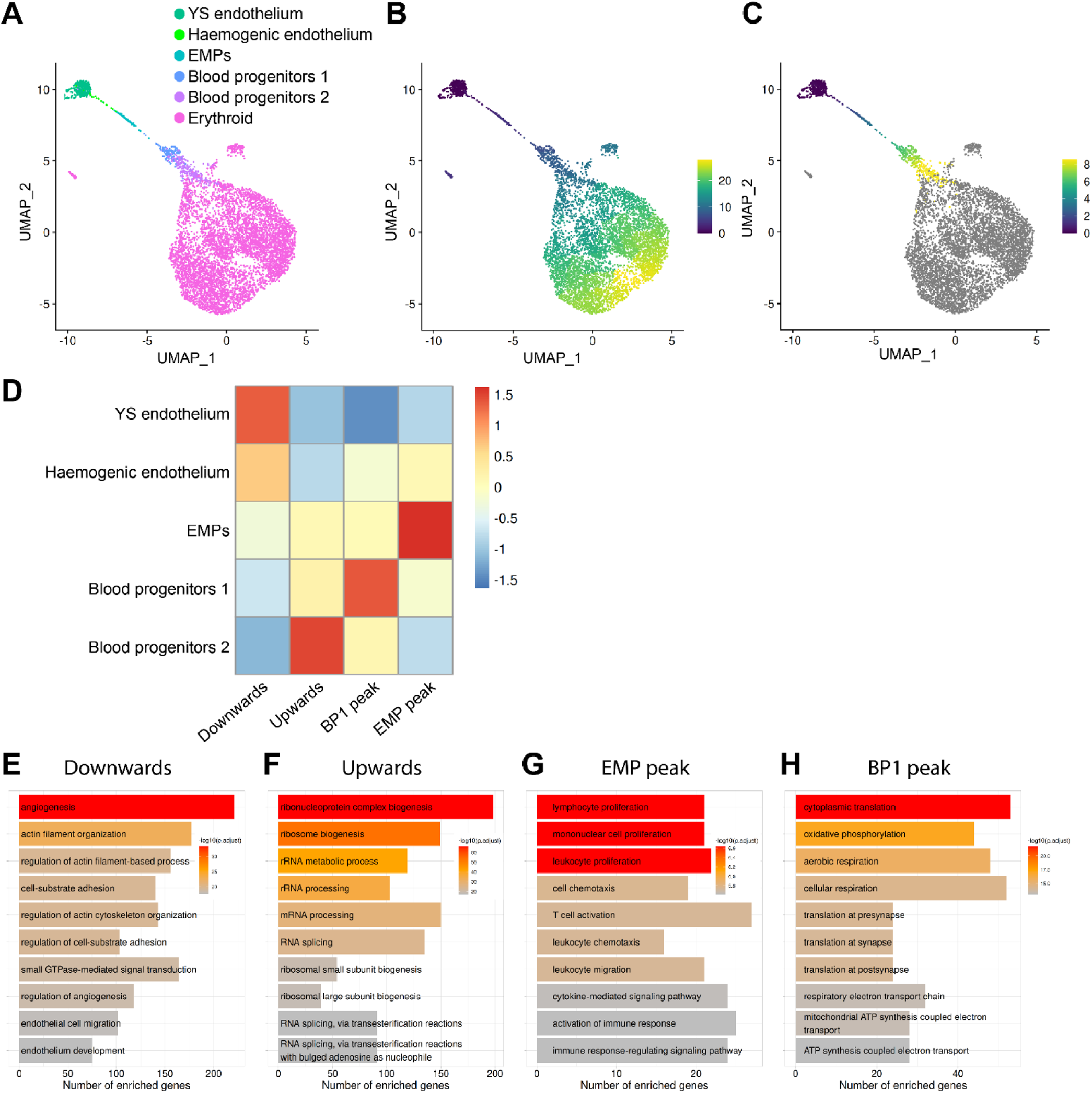
Pseudotime analysis identifies co-regulated genes and enriched GO pathways during EHT. **A** UMAP obtained after subsetting and re-clustering yolk sac haemato-vascular cells after integration of E8.5 data in E-MTAB-11763 and E-MTAB-6967, and showing our refined cluster annotation. **B,C** Pseudotime analysis highlights erythroid generation and progressive maturation **(B**) and, after excluding the erythroid cluster (**C**), the progressive differentiation from yolk sac endothelial cells to haemogenic endothelial cells and then EMPs followed by blood progenitors 1 and 2. The scale bars represent pseudotime. **D** Genes that changed as a function of pseudotime were identified and grouped according to their expression patterns. Heatmap shows 4 groups of co-regulated genes during differentiation from yolk sac endothelium to blood progenitors 2. **E-H** Gene ontology analysis for each group of co-regulated genes. YS: yolk sac; EMP: erythro-myeloid progenitor; BP1: blood progenitors 1.

Next, we took advantage of a method that analyses genes that change as a function of pseudotime (Sharon *et al*., 2019) to identify genes that showed an upward or downward trend during EHT (**Fig. 5D**, **Table S1**). We found that 2,924 genes were progressively downregulated, whereas 1,776 genes were progressively upregulated, during the predicted transition from endothelial to haematopoietic cells. We also found that 231 genes were upregulated during the predicted transition from ECs to EMPs, but downregulated during subsequent differentiation towards BP1 and then BP2 (peak transcript levels in EMPs). Moreover, 848 genes were progressively upregulated from ECs to BP1, but downregulated in BP2 (peak transcript levels in BP1) (**Fig. 5D**).

To identify potential biological processes underpinning EHT, we used gene ontology (GO) analysis on these 4 groups of co-regulated genes (**Fig. 5E-H**, **Table S2**). The GO analysis of genes with a downward expression trend during the predicted progression from endothelial to haematopoietic cells revealed enrichment in biological processes related to angiogenesis, cytoskeleton re-organisation, cell adhesion, endothelial cell migration and endothelial development (**Fig. 5E**). Genes with an upward expression trend instead were involved in mRNA processing, splicing and ribonucleoprotein biogenesis (**Fig. 5F**). Genes that peaked in EMPs showed enrichment in terms involved in leukocyte migration, myeloid proliferation and immune response (**Fig. 5G**), in agreement with EMPs being responsible for the production of foetal blood and immune cells. Finally, genes that peaked in BP1 are involved in oxidative phosphorylation, cellular respiration and ATP synthesis (**Fig. 5H**), raising the possibility that these cells have increased metabolism.

To identify genes belonging to known signalling pathways amongst each one of the 4 groups of co-regulated genes, we performed pathway analysis using the WikiPathways resource (Agrawal *et al*., 2024) (**Table S3**). Genes that were downregulated when ECs were predicted to transition to haematopoietic cells belonged to the cholesterol biosynthesis, IL3, IL17A, PI3K/AKT/mTOR and EFGR1 signalling pathways. Genes that were upregulated during this transition are involved in RNA processing and cell cycle activation. Genes peaking in BP1 are involved in the regulation of cell metabolism, whereas genes peaking in EMPs belonged to the TYROBP and IL5 signalling pathways (**Table S3**). Notably, by comparing EMPs to other yolk sac clusters, we identified multiple genes within the TYROBP signalling pathway that, in addition to peaking in EMPs, were significantly enriched in EMPs compared to the other cells in the haemato-vascular cluster (**Fig. S2**).

Overall, this analysis provided insights into molecular changes predicted to accompany EHT and haematopoietic differentiation in the yolk sac, and identified candidate signalling pathways that may be studied further to understand the molecular control of these complex cellular differentiation pathways.

### Differential expression analysis identifies EMP and haemogenic endothelium markers

To identify markers for haemogenic ECs and EMPs, we used our integrated E8.5 mouse dataset to perform differential expression (DE) analysis of endothelial and haematopoietic cell populations in both the yolk sac and embryo. First, we subset the clusters for both cell populations and then performed DE analyses by comparing clusters in different combinations to identify the top differentially regulated genes. To identify genes that may encode markers useful for cell identification via immunostaining or flow cytometric cell sorting, we filtered DE genes for those that encoded transmembrane proteins (**Table S4**).

By comparing EMPs to haematopoietic and endothelial cells, we identified *Csf1r* and *Alox5ap* as enriched in EMPs (**Fig. 6A**), agreeing with previous studies that identified them as EMP markers (Hoeffel *et al*., 2015; Sharon *et al*., 2019). In addition, we identified *Fcer1g*, *Tyrobp* and *Mctp1* as potential new markers for EMPs (**Fig. 6A,B**), whereas *Fxyd5* and *Scarf1* were identified as markers of yolk sac ECs and EMPs, and to a lower extent of BP1 (**Fig. 6B**). None of these seven genes were expressed in more differentiated BP2 and erythroid cells. A comparison of genes expressed in haemogenic ECs versus EMPs showed that acquisition of an EMP identity was associated with a loss of *Stab2* (**Fig. 6C**), as previously described (Shvartsman *et al*., 2019). Furthermore, we identified *Icam2*, *Cldn5* and *Gpr182* as additional genes that were expressed in yolk sac ECs and haemogenic ECs but downregulated in EMPs (**Fig. 6C**).

**Fig 6.**
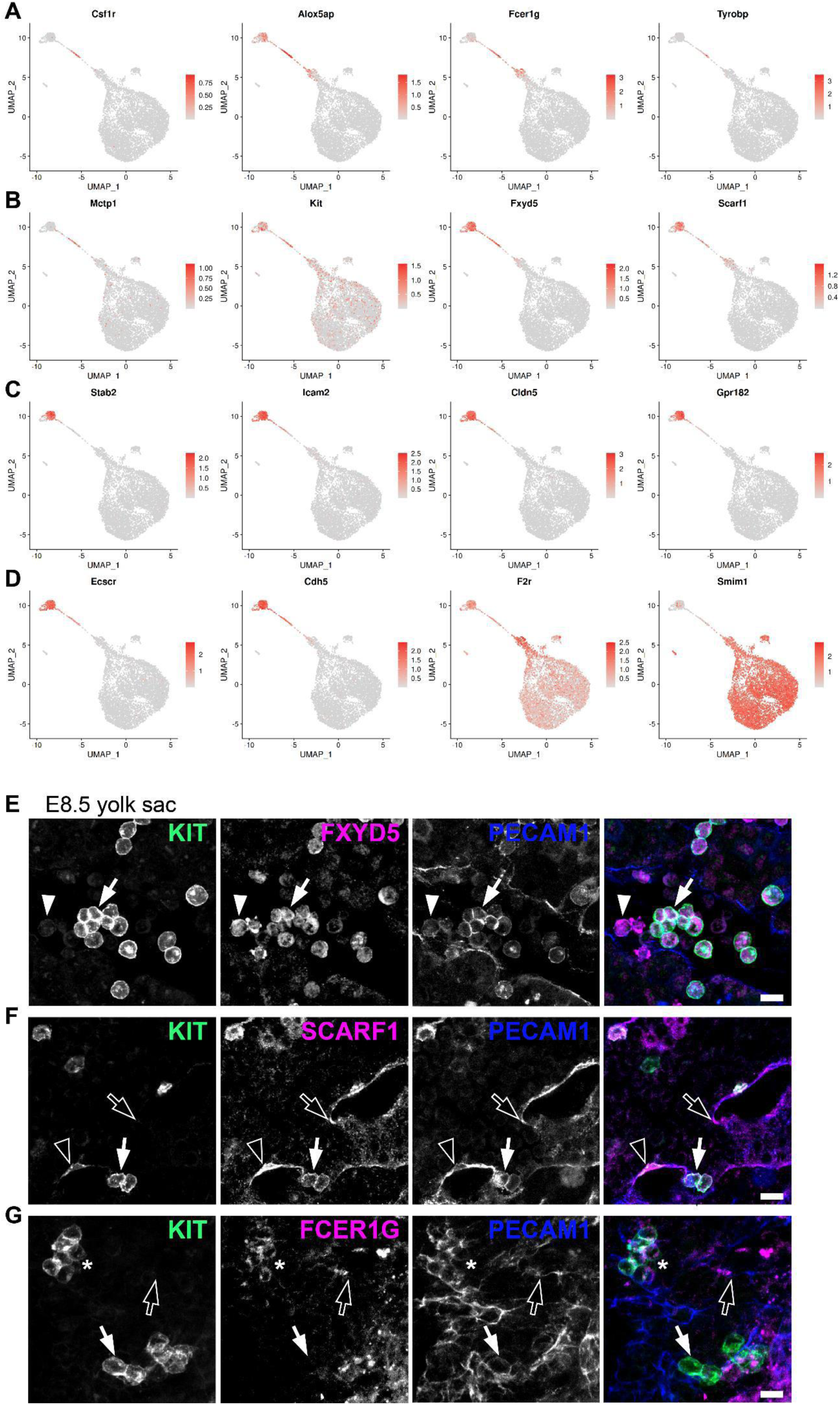
Differential gene (DE) expression analysis and immunostaining validation identifies new markers for haemogenic endothelial cells and EMPs. **A-D** UMAP representation of transcript levels for the indicated marker genes identified by DE analysis between the different yolk sac haemato-vascular clusters. The scale bars represent log-normalised transcript levels. **E-G** Immunofluorescence staining of E8.5 yolk sac for the EMP marker KIT together with PECAM1 and FXYD5, SCARF1 or FCER1G. Emerging EMPs are indicated with arrows (round KIT+ PECAM1+ cells), presumed blood (myeloid) progenitors with arrowheads (round KIT- PECAM1low cells), endothelial cells with empty arrows (flat KIT- PECAM1+ cells), haemogenic endothelial cells with empty arrowheads (flat KIT+ PECAM1+ cells) and mature EMPs with asterisk (round KIT+ PECAM1+ FCER1G+ cells). Scale bars: 50 µm; n = 3 embryos.

By comparing EMPs to BP1, and BP1 to BP2, we identified transcriptional changes associated with the EMP transition to more committed progenitor states. This transition was associated with the downregulation of the endothelial genes *Ecscr* and *Cdh5*, the EMP marker *Alox5ap*, as well as the downregulation of our predicted novel EMP marker *Fcer1g* and yolk sac EC and EMP markers *Fxyd5* and *Scarf1* (**Fig. 6A-D**). By contrast, the EMP transition to more committed progenitor states was associated with the upregulation of the megakaryocytic gene *F2r* and the erythroid gene *Smim1* (**Fig. 6D**).

To confirm that *Fxyd5*, *Scarf1* and *Fcer1g* encode EMP and haemogenic endothelial markers, we performed immunofluorescence staining of E8.5 yolk sac. Although we had found *Fxyd5* transcripts in yolk sac ECs and EMPs, FXYD5 staining was restricted to round KIT^+^ PECAM1^+^ cells consistent with newly emerging EMPs, and there was no obvious staining of ECs (**Figs. 6E, S3A**). Additionally, FXYD5 immunostaining identified round KIT^-^PECAM1^low^ cells, agreeing with our transcriptomic analysis detecting BPs with no *Kit* transcripts and low levels of *Fxyd5* (**Fig. 6B**). Consistent with the presence of *Scarf1* transcripts in both yolk sac ECs and EMPs, immunofluorescence staining identified SCARF1 protein in flat KIT^-^ PECAM1^+^ ECs and round KIT^+^ PECAM1^+^ emerging EMPs. Furthermore, SCARF1 immunostaining identified flat KIT^+^ PECAM1^+^ cells that may be haemogenic ECs (**Figs. 6F, S3B**). Although *Fcer1g* transcripts were detected in both EMPs and BP1, immunofluorescence identified FCER1G protein within a subset of round KIT^+^ cells. Additionally, FCER1G immunostaining identified flat KIT^-^ PECAM1^+^ ECs, consistent with our transcriptomic analysis detecting low levels of *Fcer1g* transcripts in yolk sac ECs (**Figs. 6G, S3C**).

A comparison with clusters from the embryo showed that the markers identified in our analysis as expressed in yolk sac ECs (*Fxyd5*, *Scarf1*, *Icam2*, *Cldn5*, *Gpr182*) were also detected in the endocardium and in venous ECs (**Fig. 7A-C**). In contrast, the EMP markers we identified (*Csf1r*, *Alox5ap*, *Fcer1g*, *Tyrobp*, *Mctp1*) were not expressed in ECs from the embryo, confirming their specificity. Nevertheless, these markers were also expressed in EMP-like cells from the embryo, thus corroborating their appropriate annotation.

**Fig 7.**
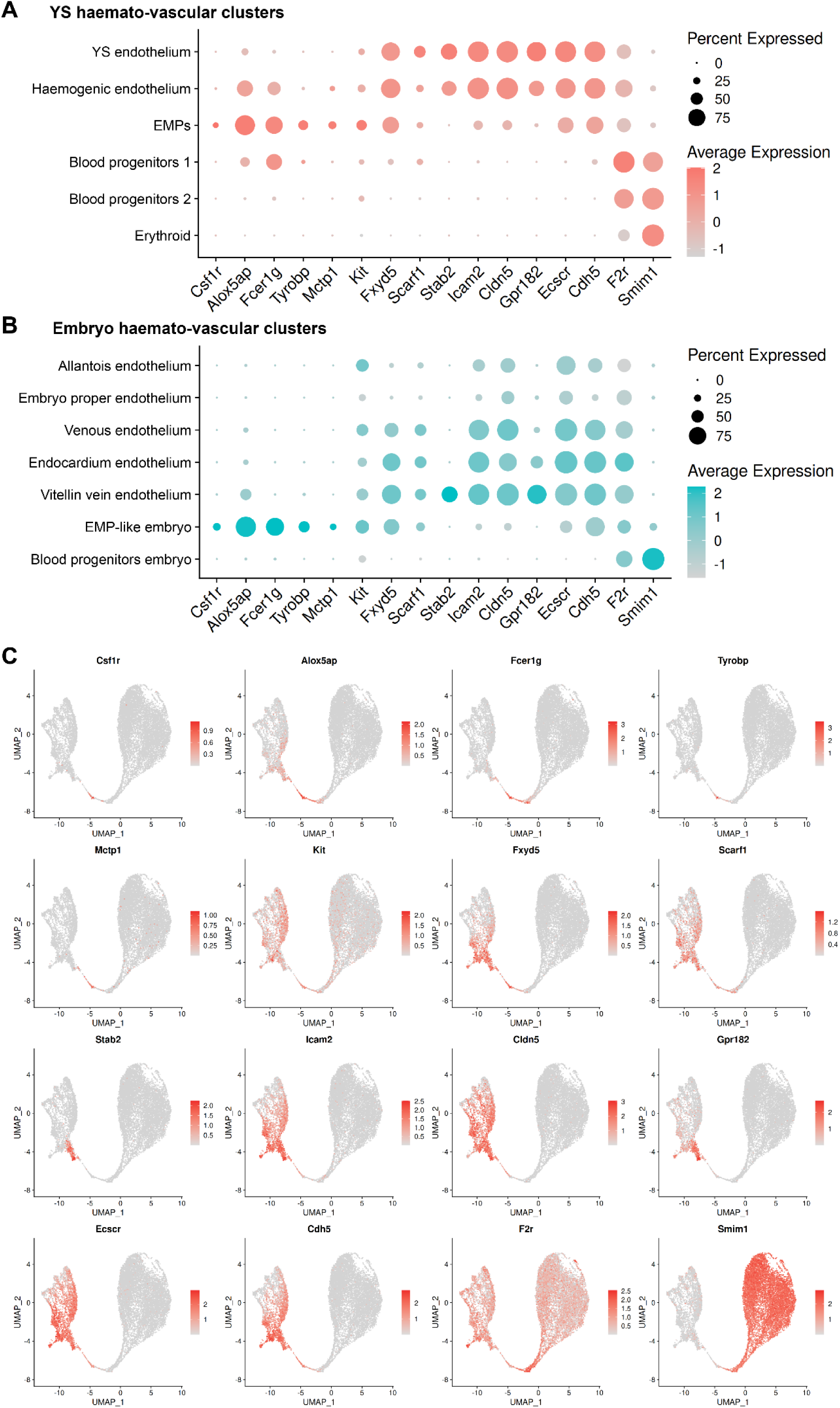
Expression analysis in haemato-vascular cells from both yolk sac and embryo shows specificity of the new markers for haemogenic ECs and EMPs. **A,B** Dot plot representation of markers identified by DE analysis in (**A**) yolk sac and (**B**) embryo haemato-vascular clusters. The dot size reflects the percentage of cells in each cluster expressing the gene. The scale bar indicates log-normalised gene expression. **C** UMAP of the E8.5 haemato-vascular clusters from both embryo and yolk sac showing transcript levels for the indicated markers identified by DE analysis. The scale bars represent log-normalised transcript levels.

Taken together, these findings show that novel EMP markers can be identified by scRNAseq analysis and that they could be used to investigate both *bona fide* EMPs in the yolk sac as well as poorly characterised EMP-like cells found in the embryo.

## DISCUSSION

To capture the time when EMPs emerge from the yolk sac haemogenic endothelium and to define the transcriptome of the cell populations involved in EHT, we re-analysed transcriptomic data from the E8.5 mouse embryo (**Fig. 1**). Our analysis provided a refined annotation of endothelial and haematopoietic populations in the mouse yolk sac (**Fig. 3A,B**) and identified a subset of ECs co-expressing endothelial and haematopoietic markers, consistent with a haemogenic endothelial phenotype (**Fig. 4A,B**). This refined annotation also defined E8.5 yolk sac EMPs and demonstrated co-expression of myeloid, erythroid and progenitor markers in this cluster, consistent with the known dual erythro-myeloid differentiation potential of EMPs. Moreover, we identified EMP-like cells within the E8.5 embryo proper, which have not been highlighted in prior transcriptomic studies of early haematopoiesis (**Fig. 3B**). Knowing that haematopoietic cells enter the embryo only from E9.5, when the heart begins to beat to initiate blood circulation (Frame *et al*., 2016; Kasaai *et al*., 2017), we considered two possibilities to explain the existence of EMP-like cells in the embryo. On the one hand, these cells may represent yolk sac-derived haematopoietic progenitors that had contaminated the embryonic samples during dissection. On the other hand, it is conceivable that these cells represent EMP-like cells originating in the embryo rather than in the yolk sac. The latter possibility is consistent with our recent discovery of a *Pax3* lineage-derived EMP-like progenitor in the mouse embryo (Canu *et al*., 2024). However, further work is required to accurately describe these EMP-like cells.

Others have used scRNAseq to map potential differentiation trajectories for EMPs in the yolk sac (Zhao *et al*., 2023). Additionally, our analysis identified distinct BP1 and BP2 clusters as consecutive stages of EMP maturation (**Fig. 5A-C**) and provided new insight into candidate signalling pathways controlling this transition (**Fig. 5D-H**). As primitive erythrocytes are produced from E7.0 onwards (Palis *et al*., 1999; Ferkowicz and Yoder, 2005), but EMPs only emerge from E8.5 onwards (Frame, McGrath and Palis, 2013; Gomez Perdiguero *et al*., 2014; Hoeffel *et al*., 2015; McGrath *et al*., 2015), the erythroid cluster in the E8.5 dataset is expected to contain mostly primitive erythrocytes that were produced prior to or alongside EMP emergence, with only a potential minor contribution from EMP-derived erythrocytes. No markers of fully differentiated macrophages or other myeloid cell types were observed, confirming previous studies showing that macrophages form in the yolk sac from E9.0 onwards (Hoeffel and Ginhoux, 2018). We also did not detect a lymphoid cluster at E8.5, in agreement with prior reports of LMP presence in the yolk sac only from E9.5 onwards (Böiers *et al*., 2013).

Previous studies established that EHT is associated with downregulation of endothelial genes and concomitant upregulation of haematopoietic genes (Palis *et al*., 1999; Frame, McGrath and Palis, 2013; Gritz and Hirschi, 2016; Lilly *et al*., 2016; Canu *et al*., 2020). Here, pseudotime reconstruction highlighted dynamic gene expression trends during EHT and haematopoietic maturation in the yolk sac (**Fig. 5A-D**), whereby haemogenic ECs form a transcriptional continuum bridging the transition from endothelial to haematopoietic cell identities, thus implying a gradual and continuous process rather than a discrete lineage switch. Altogether, these findings align with established models of yolk sac haematopoiesis derived from genetic lineage tracing, flow cytometry experiments and prior transcriptomic analysis (Frame, McGrath and Palis, 2013; Hoeffel and Ginhoux, 2018; Canu and Ruhrberg, 2021; Sánchez-Lanzas, Jiménez-Pompa and Ganuza, 2024; Sugimura *et al*., 2025).

The progressive downregulation of genes associated with angiogenesis, cytoskeletal organisation and endothelial adhesion during the predicted EHT (**Fig. 5E**) may indicate structural and functional changes necessary for cell detachment and transition to a haematopoietic phenotype. Conversely, the upregulation of genes associated with RNA processing (**Fig. 5F**) suggests that haematopoietic specification is coupled to enhanced transcriptional activity. Furthermore, upregulation of genes associated to cell cycle activation and transition from the G1 to S phase (**Table S3**) agrees with previous findings that EHT requires cell cycle progression and is controlled by the cell cycle regulators CDK4/6 and CDK1 (Marcelo *et al*., 2013; Canu *et al*., 2020).

DE analysis identified components of the TYROBP signalling pathway as significantly enriched in EMPs (**Fig. S2**). In the adult, the activation of the TYROBP signalling pathway has been associated with the microglial response to stress (Haure-Mirande *et al*., 2022; Zhang *et al*., 2024). However, to the best of our knowledge, this signalling pathway has not been previously associated with EMP emergence. Further investigation should determine whether this pathway may be functionally involved in the production of EMPs or other haematopoietic progenitors (*e.g.*, HSCs from the dorsal aorta) and whether manipulating TYROBP *in vitro* during differentiation of induced pluripotent stem cells may allow to devise more efficient haematopoietic differentiation methods.

DE analysis also identified upregulation of genes associated with oxidative phosphorylation upon differentiation of EMPs to BP1 (**Fig. 5H**). Previous studies of pluripotent stem cells and neuronal progenitors showed that their differentiation is associated with metabolic reprogramming, whereby differentiating progenitors switch from glycolysis to oxidative phosphorylation (Zheng *et al*., 2016; Dahan *et al*., 2019; Mahmoud, 2023). Our analysis raises the possibility that an analogous process may occur when haematopoietic progenitors in the yolk sac commit to differentiation.

Our DE analysis identified both known and previously undescribed candidate markers associated with the formation of haemogenic endothelium and EMPs (**Fig. 6A-D**). Established EMP markers such as *Csf1r* and *Alox5ap* were previously described by others (Hoeffel *et al*., 2015; Sharon *et al*., 2019), thus validating the robustness of our transcriptomic analysis. In addition, we identified *Fcer1g*, *Tyrobp*, and *Mctp1* as potential novel EMP markers, and *Fxyd5* and *Scarf1* as shared markers between yolk sac endothelial and haematopoietic cells. The expression of *Fxyd5* and *Scarf1* in endothelial and EMP clusters, and to a lower extent in BP1, but not in more differentiated cells, suggests that these genes may serve as new markers for yolk sac ECs and emerging EMPs. Immunofluorescence staining of E8.5 yolk sac corroborated the predictions of our DE analysis: FXYD5 was present on round KIT^+^ PECAM1^+^ cells consistent with emerging EMPs (**Figs. 6E, S3A**); SCARF1 labelled flat KIT^-^ PECAM1^+^ ECs, flat KIT^+^ PECAM1^+^ cells consistent with haemogenic ECs and round KIT^+^ PECAM1^+^ cells consistent with emerging EMPs (**Figs. 6F, S3B**); FCER1G marked flat KIT^-^ PECAM1^+^ ECs and a few round KIT^+^ PECAM1^+^ cells that may represent a subset of EMPs (**Figs. 6G, S3C**). Nevertheless, these experiments also highlighted differences in RNA and protein expression. Thus, although *Fxyd5* RNA was detected in yolk sac ECs, haemogenic ECs and EMPs, and at lower levels in BP1, the FXYD5 protein appeared to be prevalent in round KIT^+^ PECAM1^+^ cells consistent with EMPs and in round KIT^-^ PECAM1^low^ cells presumed to be myeloid-committed BP1 (**Figs. 6E, S3A**). While *Fcer1g* RNA clearly identified the EMP cluster, the FCER1G protein was detected in only a subset of round KIT^+^ PECAM1^+^ cells, possibly marking more mature EMPs (**Figs. 6G, S3C**). This is reminiscent of observations for *Csf1r*, a gene expressed in EMPs at the RNA level, whereas CSF1R protein only becomes translated upon differentiation to myeloid cells (Hoeffel *et al*., 2015). These discrepancies between RNA and protein expression may be due differences in expression dynamics and half-lives for mRNAs and proteins (Vogel and Marcotte, 2012). Overall, these immunostaining results validated our transcriptomic predictions and highlight the utility of single-cell datasets in discovering functional cell surface markers. Previously used markers for haemogenic ECs include the surface markers CDH5, KIT and CD41, and haematopoietic transcription factors such as RUNX1 (Mikkola *et al*., 2003; Nadin, Goodell and Hirschi, 2003; Lilly *et al*., 2016), whereas previously used markers for EMPs include KIT, CD41 and CD16/32, as well as the gene expression reporter *Csf1r*-eGFP (McGrath *et al*., 2011; Azzoni *et al*., 2018; Plein *et al*., 2018). The addition of the novel markers identified in our analysis will provide complementary tools to define haemogenic ECs and EMPs in the mouse yolk sac.

In summary, this study refines the cellular and molecular characterisation of yolk sac haematopoiesis, provides a detailed transcriptomic roadmap of EHT in the yolk sac and identifies novel candidate markers for haemogenic endothelium and EMPs. The identification of FXYD5, SCARF1 and FCER1G as markers for EMPs and haemogenic ECs may offers valuable tools for future *in vivo* and *in vitro* studies. Finally, future studies may wish to determine whether the novel EMP markers are present also in foetal HSCs, and whether they are conserved in human haemopoiesis. Defining the specificity of such novel markers may allow researchers to better define EMP identity and lineage potential as a prerequisite for deciphering the ontogeny of the immune system and, ultimately, to harness early haematopoietic pathways for regenerative medicine.

## ACKNOWLEDGEMENTS

We thank the staff of the Biological Resources, FACS and Imaging Facilities at the UCL Institute of Ophthalmology for their technical support. This research was supported by grants from Wellcome (205099/Z/16/Z) and the British Heart Foundation (PG23/11301).

## AUTHORSHIP CONTRIBUTIONS

GC, GDP and CR conceived and designed the study. GC and CR co-wrote the manuscript. GC and AM performed immunostaining experiments. GDP performed bioinformatic analyses. GC and CR supervised the project. All authors read and approved the submitted manuscript.

---

**Fig S1.**
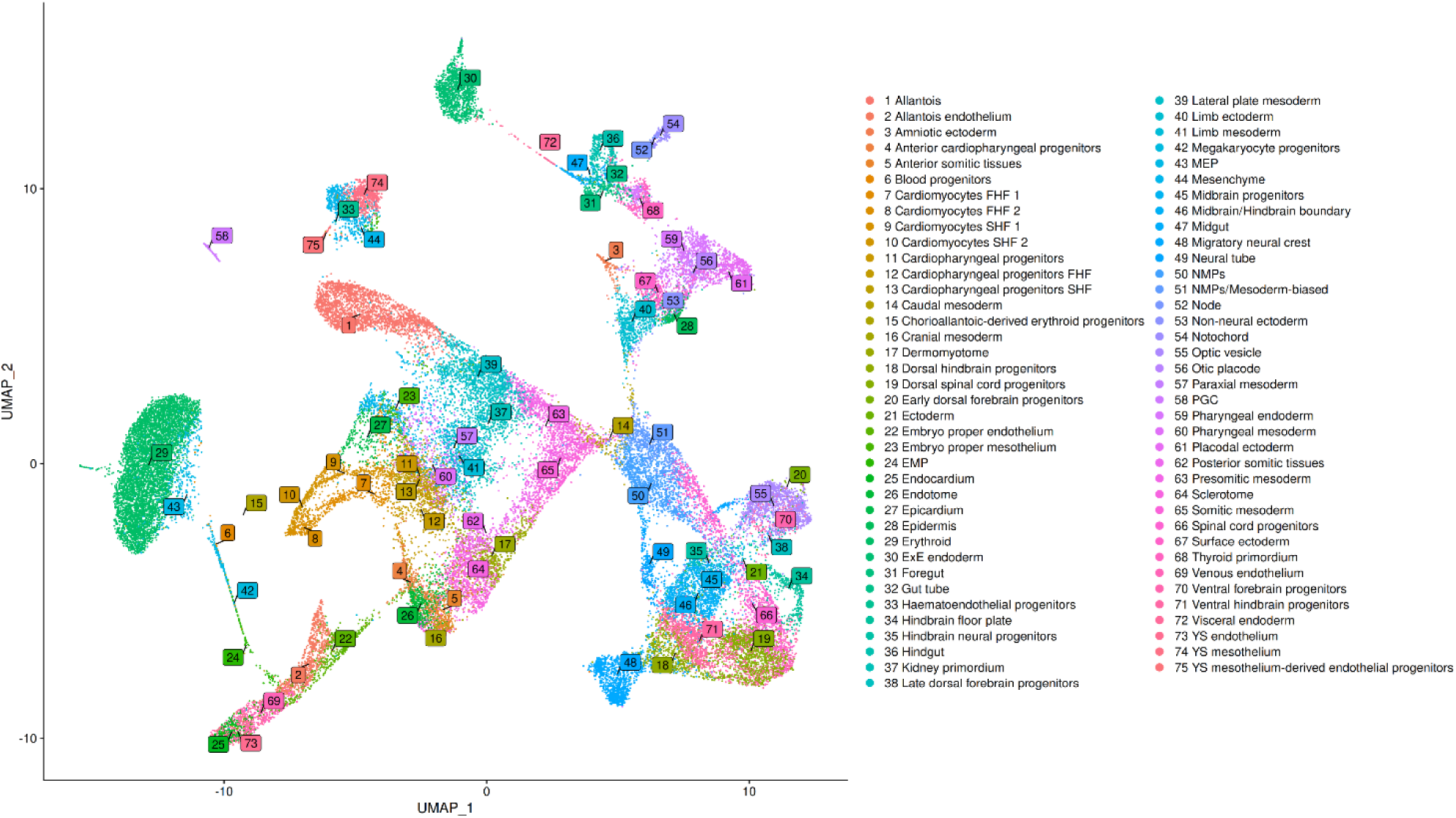
Cluster annotation of E8.5 mouse scRNAseq data in E-MTAB-11763. UMAP of the E8.5 cells in E-MTAB-11763 dataset, labelled with the cluster annotations provided by the authors of the original publication (Goh et al., 2023).

**Fig S2.**
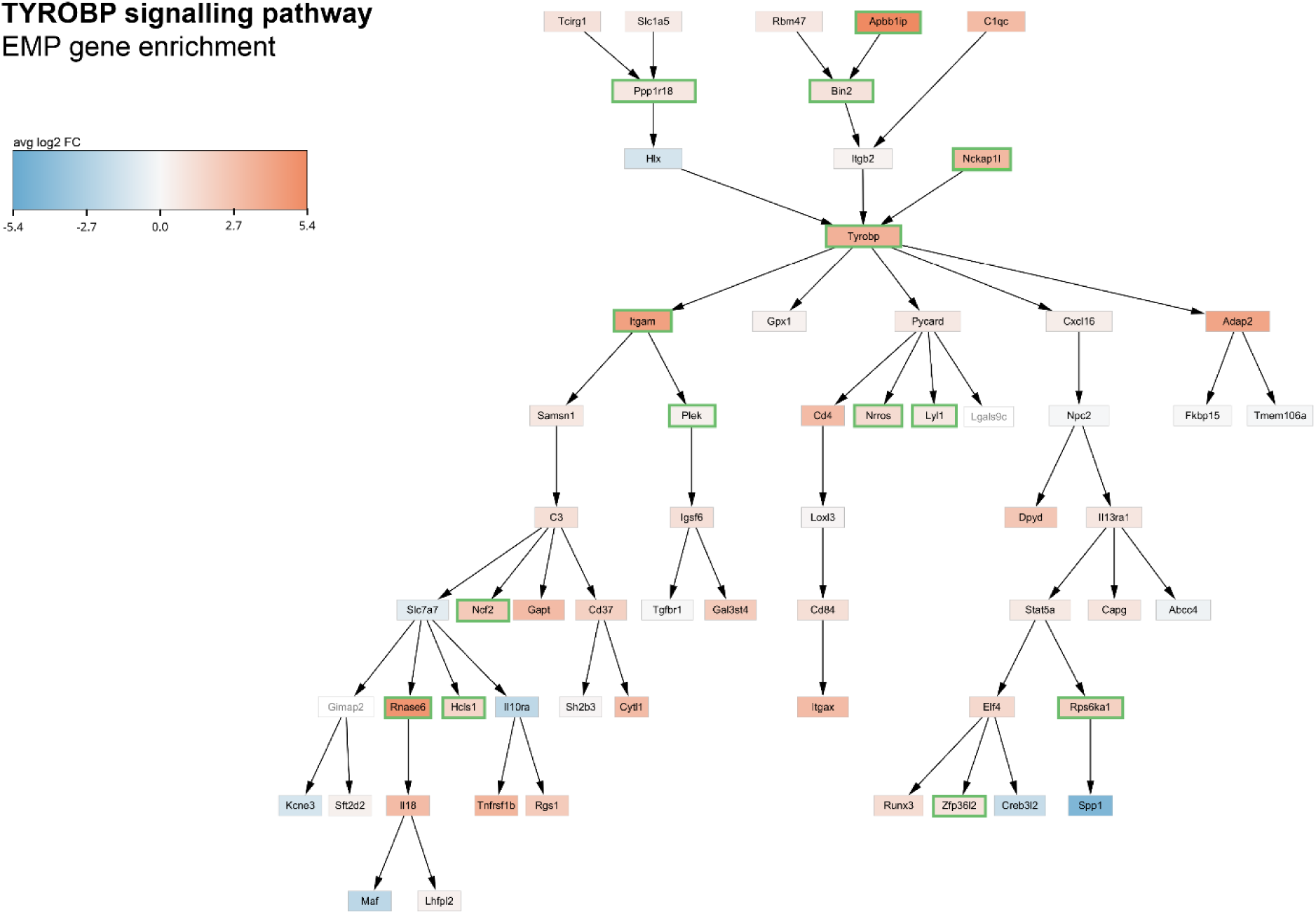
The TYROBP signalling pathway is enriched in emerging EMPs. TYROBP pathway genes enriched in EMPs compared to haemato-vascular clusters from both yolk sac and embryo. The box colour indicates the average log2 fold change. A green border indicates significant enrichment in EMPs.

**Fig S3.**
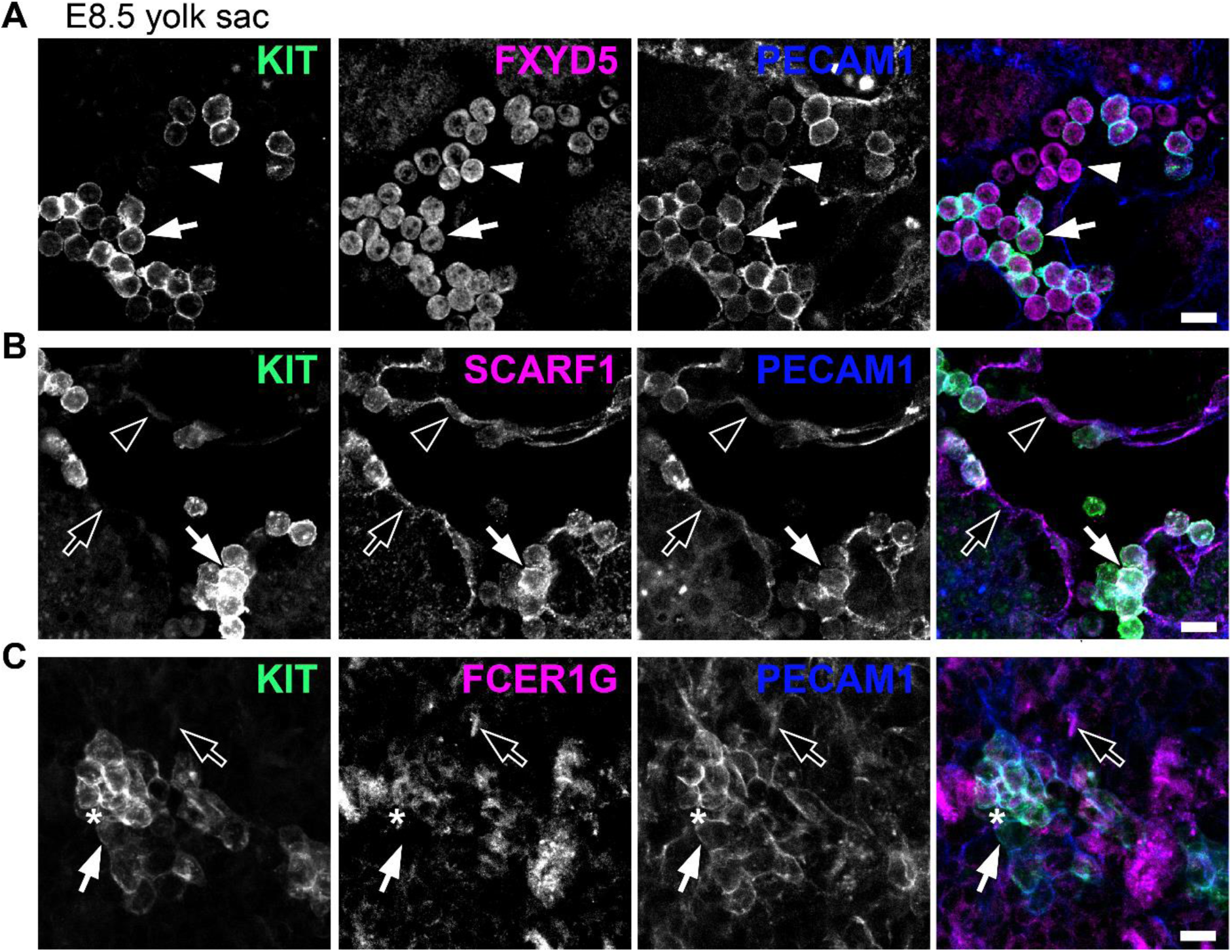
Immunostaining validation of new markers for haemogenic endothelial cells and EMPs. **A-C** Immunofluorescence staining of E8.5 yolk sac for the EMP marker KIT together with PECAM1 and FXYD5, SCARF1 or FCER1G. Emerging EMPs are indicated with arrows (round KIT+ PECAM1+ cells), presumed blood (myeloid) progenitors with arrowheads (round KIT- PECAM1low cells), endothelial cells with empty arrows (flat KIT- PECAM1+ cells), haemogenic endothelial cells with empty arrowheads (flat KIT+ PECAM1+ cells) and mature EMPs with asterisk (round KIT+ PECAM1+ FCER1G+ cells). Scale bars: 50 µm; n = 3 embryos.

